# Incomplete reprogramming of cell-specific epigenetic marks during asexual reproduction leads to heritable phenotypic variation in plants

**DOI:** 10.1101/267955

**Authors:** Anjar Wibowo, Claude Becker, Julius Durr, Jonathan Price, Stijn Spaepen, Sally Hilton, Hadi Putra, Ranjith Papareddy, Quentin Saintain, Sarah Harvey, Gary D. Bending, Paul Schulze-Lefert, Detlef Weigel, Jose Gutierrez-Marcos

## Abstract

Plants differ from animals in their capability to easily regenerate fertile adult individuals from terminally differentiated cells [1]. This unique developmental plasticity is commonly observed in nature where many species can reproduce asexually through the ectopic initiation of organogenic or embryogenic developmental programs [2, 3]. However, it is not currently known if this developmental reprogramming is coupled to a global epigenomic resetting, or what impact it has on the phenotype of the clonal progeny. Here we show that plants asexually propagated via induction of a zygotic developmental program do not fully reset cell-specific epigenetic imprints. These imprints are instead inherited even over multiple rounds of sexual reproduction, becoming fixed in hybrids and resulting in heritable molecular and physiological phenotypes that depend on the founder cell used. Our results demonstrate how novel phenotypic variation in plants can be unlocked through the incomplete reprogramming of cell-specific epigenetic marks during asexual propagation.

## Main Text

Compared to animals, somatic cells of plants can be much more easily coaxed into regenerating entire individuals. Asexual reproduction is therefore much more common in plants than in animals, and this has been traditionally exploited by humans for the clonal propagation and genetic manipulation of many economically important plant species [4]. Although clonal propagation provides ecological and evolutionary benefits, the resulting restricted genetic variation could be detrimental to fitness [5, 6]. Notably, clonally propagated plants are not always phenotypically identical to their parents; a phenomenon often attributed to the accumulation of genetic mutations. Yet there is little direct evidence that genetic changes are solely responsible for this phenotypic variation, thus we hypothesized that the phenotypic diversity apparent in clonal plants may have epigenetic underpinnings. Because plants can reproduce asexually from below-ground and above-ground organs, which are known to be epigenetically distinct [7, 8] we took advantage of this situation to determine to what extent the epigenome could influence clonal plant phenotypes. Specifically, we created somatic embryos from distinct root (Root Origin, RO) and leaf (Leaf Origin, RO) tissues of *Arabidopsis thaliana* (Col-0 strain) by the controlled expression of a RWP-RK zygotic factor [9] (Supplementary Fig. 1), in order to mimic naturally-occurring events associated with asexual propagation [2, 10]. We collected seeds from independently regenerated G_0_ individuals after self-pollination and further propagated each line by selfing for over three consecutive generations (G1-G3) (Fig. 1a). Visual examination revealed no obvious morphological differences between RO and LO plants. To determine any potential differences at the molecular level, we performed whole-genome transcriptome analyses in five randomly selected G_2_ lines. Only 13 differentially expressed genes (DEGs; FDR < 0.05) distinguished roots of LO and RO plants, but almost twenty-fold more DEGs (239, FDR < 0.01) were identified when comparing leaves of RO and LO plants (Supplementary Fig. 2 and Table 1). Gene ontology (GO) analysis revealed that these DEGs were enriched for stress and defence related genes (FDR <0.5) (Fig. 1b). The genes in these functional categories (51 total) were primarily upregulated in leaves of RO plants and involved in either transcriptional regulation (22%) or cellular signaling (27%). Clustering of all samples based on DEG expression levels suggested that leaves of RO plants had partial root characteristics, sharing apparent similarity with roots from both RO and LO plants (Fig. 1c); in contrast, leaves of LO plants formed a single distinct group (Fig. 1c).

**Figure 1.**
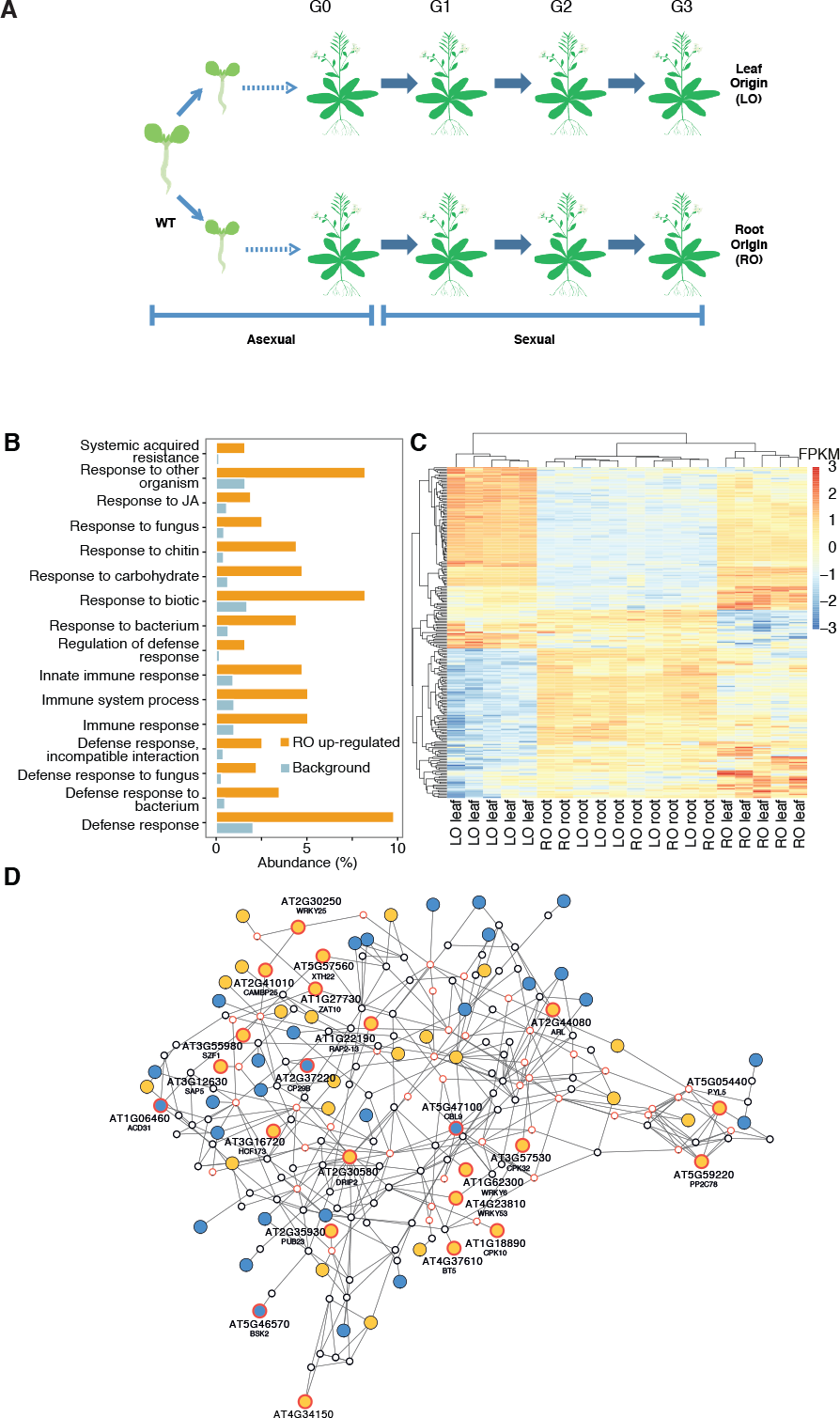
Plants regenerated from leaves and roots differ in activity of a discrete defense-related transcriptional network. **A)** Schematic diagram describing the experimental design for assessing the impact of regeneration through somatic embryogenesis from roots (RO) and leaves (LO). Regenerated plants (n=5 independent lines each) were propagated through self-fertilization over three generations to determine stability of acquired traits. **B)** Gene ontology analysis of differentially expressed genes (DEGs) between leaves of RO and LO plants reveals enrichment (FDR < 0.01) for defense-related functions. **C)** Heatmap of scaled, normalized log-transformed read counts (FPKM, Fragments Per Kilobase per Million) for genes underlying the significant enrichment of GO terms in (A). **D)** Interaction gene network of DEGs (FDR < 0.01). Of 4,585 DEGs, 2,752 were represented in the ANAP database. Nodes represent genes; triangles highlight defense-related genes from (A); edges indicate evidence for gene interaction. Blue filled circles, high expression in LO; Orange filled circles, high expression in RO; Red outlined circles, genes with stress related GO terms.

To gain further insight into this transcriptional variation, we performed a network analysis of the DEGs [II]. Out of the 239 DEGs that distinguished leaves of RO and LO plants (Supplementary Fig. 1 and Table 1), 213 had known interactions with other DEGs, of which 69 were part of a single functional network (Fig. 1d). This network included several genes (*WRKY6*, *SZF1*, *PYL5*, *PUB23* and *DRIP2*) that had been previously implicated in the negative regulation of abiotic and biotic stress responses [12–16].

We next sought to define whether the gene expression differences between LO and RO plants were functionally meaningful and affected whole-plant or tissue-specific functions such as interaction with microorganisms. As the latter was an obvious and easily scorable physiological function to assess, we grew plants in natural soils, and after four weeks assessed the bacterial communities that became associated with these plants. Bacterial communities from roots of RO plants differed to those from both non-regenerated plants (ANOSIM, r=0.485, p=0.007) and LO plants (ANOSIM, r=0.216, p=0.056) (Fig. 2a and Supplementary Fig. 2). In addition, the *in vitro* root colonization by *Bacillus amyloliquefaciens* FZB42, a soil-borne plant-growth-promoting bacterium [17], differed significantly between RO and non-regenerated plants (Wilcoxon Rank Sum test, W = 3862.5, p-value = 5.9e-05) (Supplementary Fig. 3). Finally, we inoculated roots of regenerated and non-regenerated plants with synthetic communities (SynComs) consisting of abundant soil- and root-derived bacterial isolates [18]. Again, the root bacterial communities differed between non-regenerated and regenerated individuals (7.3% variance explained by genotype, permutation-based ANOVA test, p=0.17; Fig. S4); this was especially obvious for the Alcaligenaceae family (Beta-proteobacteria) (Fig. 2b and Supplementary Fig. 4). Similarly, when analyzing leaves of SynCom-inoculated plants (9.6% variance explained by genotype, P=0.023), the most notable differences were observed for Xanthomonadaceae (Gammaproteobacteria) (ANOVA, p < 0.05) (Supplementary Fig. 4,5), which include several phytopathogenic strains. These divergences led us to further assess the response to known pathogens. For this, we inoculated leaves with the bacterium *Pseudomonas syringae* pv. *tomato* strain DC3000 and the oomycete *Hyaloperonospora arabidopsidis* isolate NoksI, as the *Arabidopsis thaliana* Col-0 strain lacks gene-for-gene resistance to both of these pathogens [19, 20]. We found that RO plants were more sensitive to infection by either pathogen than LO individuals, and that these differences were stably inherited for at least three generations (Fig. 2c,d). This result was in line with our previous observation of the upregulation of negative regulators of biotic stress in RO plants (Fig. 1 d).

**Figure 2.**
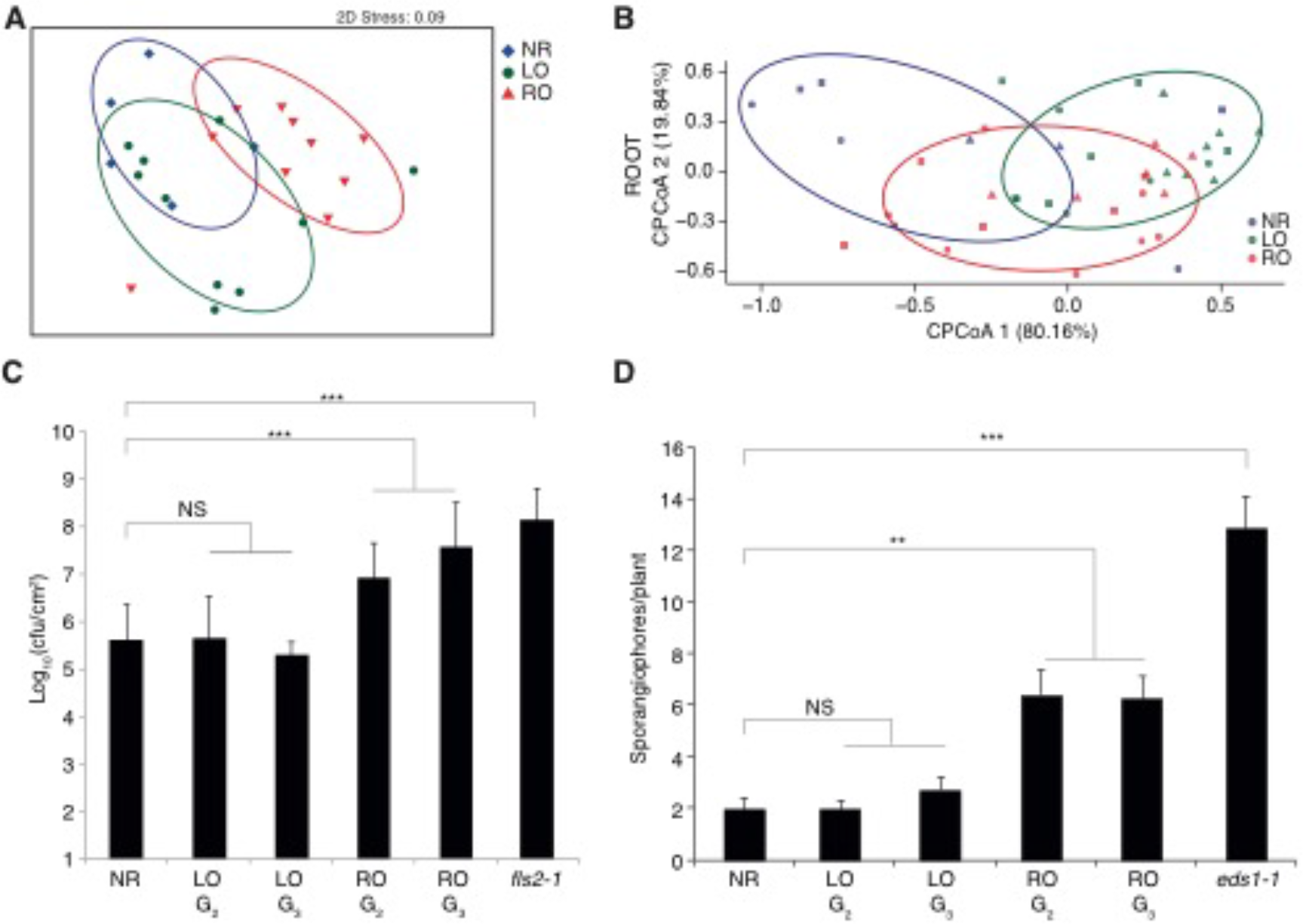
Plants regenerated from roots and leaves interact differently with microbes. **A)** PCA of Bray-Curtis distances of bacterial communities present in roots of non-regenerated and regenerated plants grown in natural soils (n=10). **B)** Canonical analysis of principal coordinates (based on Bray-Curtis distances) reveals that root- (top) and leaf- (bottom) associated bacterial communities of SynCom-colonized non-regenerated and regenerated plants differ (n=12). **D)** Susceptibility of non-regenerated and regenerated plants to *Pseudomonas syringae* pv. tomato strain DC3000 infection. Bacterial growth was determined 3 d after inoculation (100 cfu ml^-1^). Data are the means ± SD from three independent experiments. Statistical significance according to Fisher’s Exact test (*** p<0.001; NS, not significant) **E)** Susceptibility of non-regenerated and regenerated plants to *Hyaloperonospora arabidopsidis* (*Hpa*) Noks1 infection, as indicated by number of sporangiospores on leaves 3 d after inoculation (300,000 spores/ml). Data are the means from two independent experiments. The p-value was determined using Student’s *t*-test.

Given the non-expected, heritable transcriptional and phenotypic differences between RO and LO plants, which ought to be genetically identical, we investigated any potential epigenomic basis for these differences. DNA methylation is an important epigenetic mark, for which there are excellent statistical methods for genome-wide comparisons [21], and the resetting of DNA methylation during sexual reproduction has been well documented [22]. We therefore used whole-genome bisulfite sequencing (WGBS) of leaves and roots from LO and RO individuals to monitor any methylome changes over three consecutive generations

(Supplementary Table 2). Principal component analysis (PCA) of 736,413 differentially methylated positions (DMPs) discovered in pairwise contrasts revealed clear differences between root and leaf samples from non-regenerated and regenerated plants (PC1, Fig. 3a). Regenerated samples clustered according to their tissue of origin prior to regeneration and not, as one might expect, by tissue identity at the time of DNA extraction (PC2, Fig. 3a). When repeating PCA with the methylation level of positions not classified as DMPs, we found little residual variance, thus indicating a low number of false negatives (Supplementary Fig. 6).

**Fig. 3.**
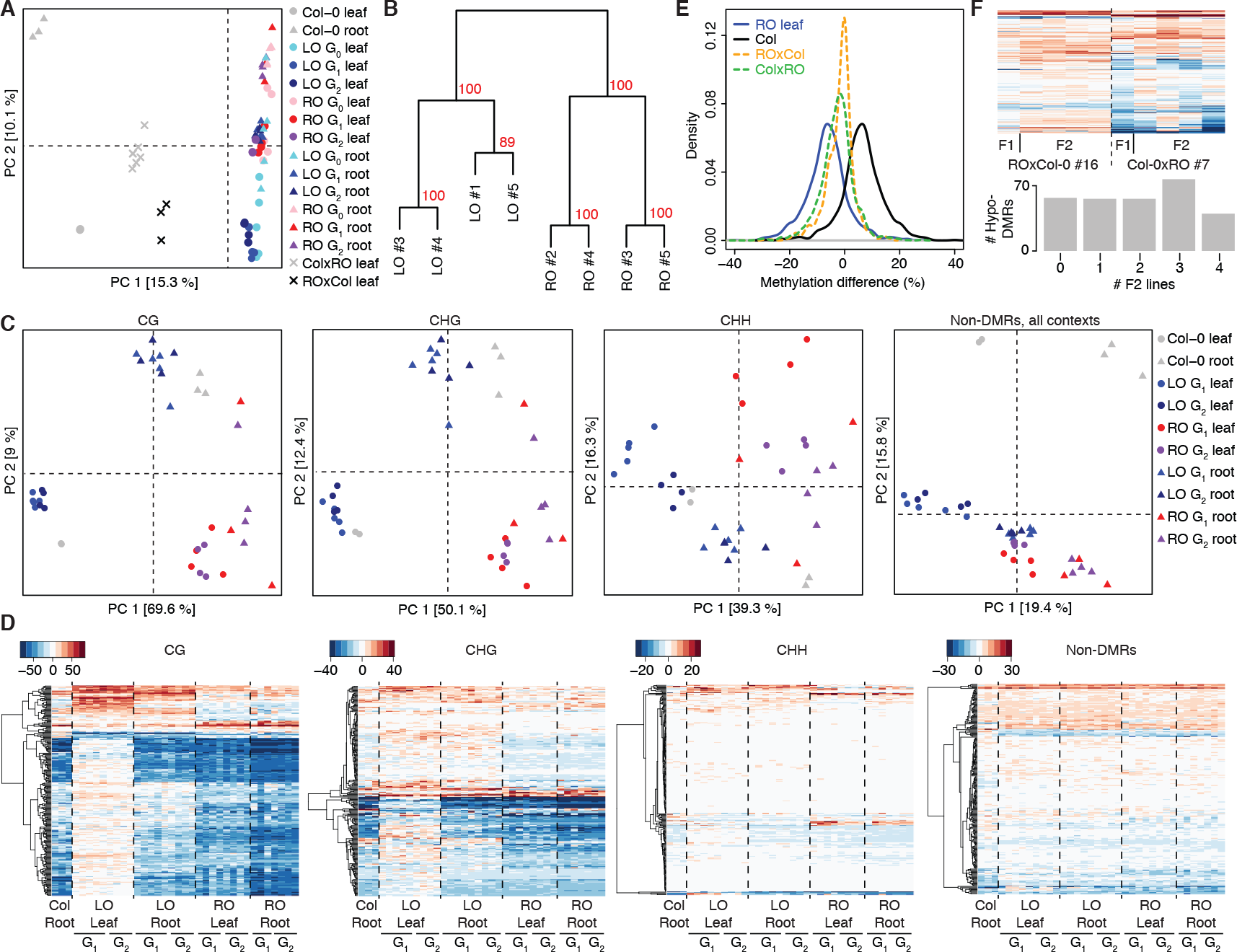
Whole-genome DNA methylation analysis reveals evidence for stable epigenetic variation in regenerated plants. **A)** Principal component analysis (PCA) of DNA methylation levels at differentially methylated positions (DMPs) identified from pairwise sample comparisons. Numbers in brackets indicate the fraction of overall variance explained by the respective PC. **B)** Clustering of LO and RO leaf samples in generation G_2_, based on 765 differentially methylated regions (DMRs) identified in all-against-all pairwise comparisons. **C)** PCA of methylation at 255 DMR loci identified in G_2_ RO vs. LO leaf comparison, divided by cytosine sequence context. Rightmost panel shows PCA on methylation in all contexts within randomly chosen non-differentially methylated regions (non-DMRs). Numbers in brackets indicate the amount of variance explained by the respective PC. **D)** Gains and losses of DNA methylation in DMRs identified between RO and LO leaves in the G_2_ generation. Color keys indicate methylation rate differences in relation to leaves of non-regenerated plants. Right-most panel shows differences in a random subset of non-DMRs. **E)** Methylation frequencies at DMR loci in leaves of non-regenerated Col-0, RO, and reciprocal crosses (F_1_) between non-regenerated Col-0 and RO plants. **F)** Methylation analysis of progeny from F_1_ reciprocal hybrids. Heatmap shows DMR methylation levels in individual F_1_ hybrid plants (#7 and #16) and each of four independent descendants. Bar plot shows frequency of hypo-methylation in F_2_ plants of DMRs that were hypo-methylated in the F_1_ hybrid.

Because the functional relevance of individual DMPs in plants is unclear, we also analyzed differentially methylated regions (DMRs). Cluster analysis based on 765 DMRs identified in pairwise comparisons between all G_2_ leaf samples validated the use of the independent LO and RO lines as two replicate groups (Fig. 3b). We found 255 consistent LO-vs-RO DMRs in G_2_ leaves (Supplementary Table 3). Methylation in LO leaves was similar to that of non-regenerated leaves (PC 1 in Fig. 3c); likewise, roots from RO samples showed patterns similar to those of non-regenerated roots. However, leaves from RO populations maintained methylation patterns that were more similar to roots, especially in symmetric cytosine contexts (Fig. 3c). Conversely, methylation levels at these DMRs in roots of LO plants were similar to those found in non-regenerated root samples (Fig. 3c), in line with previous studies in *A. thaliana* and two closely related species [7, 8]. By contrast, methylation levels at non-differentially methylated regions (non-DMRs) grouped samples primarily by their tissue identity, regardless of regenerant origin (Fig. 3c). Our DMR analysis also revealed that leaves from RO plants had less overall CG and CHG (but not CHH) methylation in these regions when compared to leaves of non-regenerated plants or LO plants (Fig. 3d). To account for sampling bias and stochastic effects, we repeated the analyses using randomly selected non-DMRs, which did not produce any evidence for reduced methylation in roots of root-derived plants (Fig. 3d). The reduced DNA methylation in RO leaves was stably inherited over at least three generations (Fig. 3c,d), indicating that root-specific DNA methylation patterns are not efficiently reset during subsequent sexual reproduction. To understand the importance of the methylome status at the time of regeneration, we compared the leaf methylome of RO plants with the methylation profiles of roots from non-regenerated seedlings and mature plants (Supplementary Fig. 7). The RO leaf methylome was more similar to that of roots from non-regenerated seedlings, implying that regeneration had been induced at the seedling stage and that the cell-specific methylation pattern was maintained throughout the regeneration process and in subsequent sexually reproduced progeny.

Because there is evidence of methylation information being transferable between chromosomes [23, 24], we tested whether such information transfer could occur *in trans* in our system. To this end, we performed reciprocal crosses between RO and non-regenerated plants. We found that in both F_1_ hybrids, DNA methylation was at mid-parent values, indicating that the DMRs on chromosomes inherited from the RO parent retained their hypomethylation status (Fig. 3e and Supplementary Fig. 8). We assessed the heritability of the observed differential methylation at these loci by sequencing individual F_2_ plant progenies. More than two-thirds (80%) of DMRs with mid-parental methylation in the F_1_ hybrid retained their hypomethylated state in at least one F_2_ descendant, indicating that allele-specific methylation was stably inherited both through mitotic and meiotic cell divisions (Fig. 3f).

Both the establishment and maintenance of DNA methylation in plants rely on a series of partially interconnected pathways, depending on the genomic features that are methylated [25]. Consequently, when genome-wide demethylation is induced by various mutations, some regions can be remethylated upon restoration of the methylation machinery, while others cannot, and this is a function of the underlying methylation pathways [26–28]. To gain insight into the mechanisms underlying the failure to reset root-specific methylation patterns in RO plants, we investigated whether methylated regions that became hypomethylated in RO plants were under the control of a specific epigenetic pathway, by comparing the methylation changes in different mutant contexts [29]. We found that CG methylation in such regions was affected in the chromatin remodelling mutant *ddm1* and in the DNA methylation maintenance mutants *met1* and *vim1 vim2 vim3*; while CHG methylation was altered in the *de novo* methyltransferase *cmt3* mutant and in mutants with a compromised H3K9 methylation machinery (Supplementary Fig. 9). Both pointed to the involvement of the RNA-directed DNA methylation (RdDM) silencing pathway, which was further supported by the hypomethylated states of these regions in *hda6* mutants, since HDA6 is an upstream component of RdDM [30]. Contrary to our expectations, the regions that were hypomethylated in RO plants were not affected in the triple *ros1 dml2 dml3* (*rdd*) mutant, which lacks three DNA demethylases, suggesting that DNA hypomethylation in RO plants is due to differences in the establishment and/or maintenance of DNA methylation in root initials during embryogenesis.

To establish a connection between the molecular and phenotypic variation generated by plant regeneration, we searched for correlations between changes in DNA methylation (604 DMRs) and gene expression (1,537 DEGs; FDR < 0.01) in leaves of regenerated plants, over two generations. We identified 29, mostly hypomethylated, DMRs that were within 2 kb up- or downstream to DEGs (Supplementary Fig. 10 and Table 4). To confirm that such DMRs indeed affect gene expression, we selected a DMR ~1 kb downstream of *RADIALIS-LIKE SANT/MYB1* (*RSM1*)/*MATERNAL EFFECT EMBRYO ARREST 3* (*MEE3*), known to regulate plant growth and flowering time [31, 32]. To assess the role of DNA methylation on this intergenic genome region we introduced an inverted repeat (IR) hairpin to direct DNA hypermethylation to the RSM1-DMR (RSM1-IR) by RdDM. Bisulfite sequencing confirmed that the targeted genome region became specifically hypermethylated in IR transgenic lines (Fig. 4a), and this was accompanied by reduced *RSM1* expression (Fig. 4b). RSM1_IR lines suffered from pleiotropic developmental defects, including accelerated senescence and earlier flowering (Fig. 4c). Transformation of RSM1_IR plants with a synthetic RSM1 construct (synRSM1) resistant to RSM1_IR targeting reversed RSM1-IR developmental phenotypes (Fig. 4d), suggesting that *RSM1* is the causative gene responsible for the underlying developmental defects observed in IR lines and that the genomic region identified acts a long-distance regulatory region influenced by DNA methylation.

**Figure 4.**
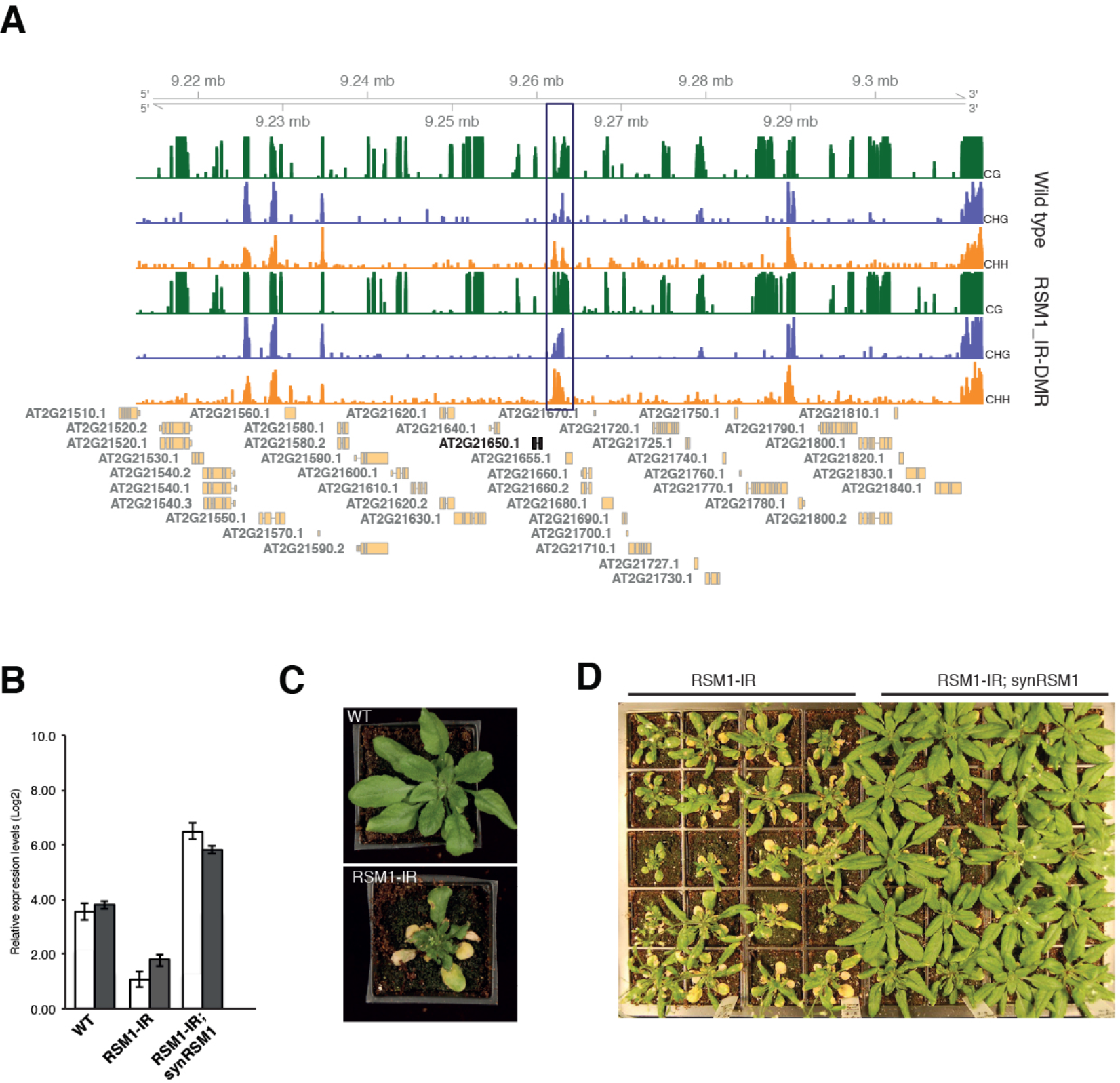
Differential methylation affects expression of *RSM1*. **A)** Snapshot of DNA methylation in different sequence contexts at the RSM/ locus in wild-type Col-0 plants and plants carrying an inverted repeat transgene (RSM1-IR). Black block highlights the region targeted by RSM1-IR-induced RdDM for DNA hypermethylation. Green bars, CG methylation; blue bars, CHG methylation; orange bars, CHH methylation. **B)** *RSM1* expression in leaves and roots of wild-type (WT), RSM1-IR plants and RSM1-IR plants complemented with synRSM 1. White bar, leaf; black bar, root. **C)** Induced DNA hypermethylation of RSM1-DMR leads to defects in plant growth and development. WT, wild-type; RSM1-IR, RSM1 inverted repeat. **D)** Genetic complementation of RSM1-IR with a synthetic RSM 1 transgene (synRSM l) resistant to RSM1-IR-induced hypermethylation.

Here, we set out to determine the cause for phenotypic variation arising from asexual propagation in plants. Our data show that, contrary to animals, plant somatic cells do not completely reset their epigenomes during cloning via the induction of an embryogenic developmental program. Because the somatic embryos induced during asexual plant reproduction closely mimics many aspects of normal embryogenesis [33], partial retention of tissue-specific epigenetic signatures would be expected in the primary regenerants, as these marks have not passed on through gametogenesis – the stage at which the epigenome is known to be reset during the plant life-cycle [22, 34]. What is surprising, however, is that progenies of regenerated plants, which have undergone multiple cycles of sexual reproduction, continue to retain cell-specific epigenetic marks, as well as transcriptional and phenotypic signatures typical of the founder tissue used for the initial propagation. The stability of these epigenetic marks resemble epialleles that are induced by the inactivation of positive regulators of DNA methylation, and which have been demonstrated to be heritable over multiple generations in epigenetic recombinant inbred lines (epiRILs) [26, 27, 35] and also exhibit stable phenotypic variation [26, 36].

Our analysis has also revealed that some epialleles captured during leaf regeneration are later reset in roots upon sexual propagation, a process that takes place during embryogenesis. The function of these imprints in roots is still unknown, but they have been found also in other plant species [7, 8, 37–39]. A tantalizing possibility is that such marks may be involved in regulating distinct tissue-specific transcriptional responses to environmental factors, such as exposure and interactions with microorganisms. In support of this argument, *A. thaliana* leaves challenged with bacterial pathogens rapidly remodel their DNA methylation profile [40] and root resistance to fungal pathogens requires active DNA demethylation [41].

Cell-specific epigenetic marks captured during somatic embryogenesis are also likely to underpin the phenotypic somaclonal variation observed in plants propagated in vitro [42–45] as well as in natural, asexually reproducing, plant populations [46]. Our findings thus not only suggest new and exciting possibilities for enhancing or, perhaps more importantly, limiting phenotypic variation in clonally propagated elite lines [44], but they also raise pertinent questions regarding the adaptive significance of the epigenomic changes captured during asexual reproduction in plants.

## Acknowledgements

We thank C. Lanz and J. Hildebrandt for help with Illumina sequencing; Rainer Borriss for providing the *B. amyloliquefaciens* FZB42 strain; and Liliana M. Costa for discussions and comments on the manuscript. Supported by ERC AdG IMMUNEMESIS, DFG SPPl529 and Max Planck Society to D.W., ERC AdG ROOTMICROBIOTA, CEPLAS and Max Planck Society to P.S.L., and BBSRC grants (BB/L003023/1, BB/N005279/1, BB/N00l94X/1 and BB/P0260lX/1) to J.G-M.

## Author contributions

G.D.B., P.S-L., D.W. and J.G-M. defined strategy, supervised and procured funding. A.W. and Q.S. performed plant regeneration, propagation and conducted genetic analyses. A.W., J.D., R.P. and S.H. performed the natural soil root microbiome analysis. J.D and S.S. performed the SynCom root and leaf microbiome analysis. J.D and Q.S. designed and performed the *B. amyloliquefaciens* root colonization assays. C.B. performed the bisulfite experiments and analyzed the data. A.W. and R.P. performed the RNAseq experiments, and J.P. analyzed the data. J.P. designed an algorithm to identify intersections between molecular datasets. A.W and J.D. created targeted hypermethylation lines and complementation lines and H.P. performed qPCR experiments. A.W., C.B., D.W. and J.G-M. wrote the paper with input from all authors.

